# Autoimmune regulator deficiency causes sterile epididymitis and impacts male fertility through disruption of inorganic physiology

**DOI:** 10.1101/2025.01.11.632558

**Authors:** Soo Hyun Ahn, Katrina Halgren, Geoffrey Grzesiak, Keith W. MacRenaris, Aaron Sue, Huirong Xie, Elena Demireva, Thomas V. O’Halloran, Margaret G. Petroff

## Abstract

Autoimmune regulator (AIRE), a transcription factor expressed by medullary thymic epithelial cells, is required for shaping the self-antigen tolerant T cell receptor repertoire. Humans with mutations in *AIRE* suffer from Autoimmune Polyglandular Syndrome Type 1 (APS-1). Among many symptoms, men with APS-1 commonly experience testicular insufficiency and infertility, but the mechanisms causing infertility are unknown. Using an *Aire*-deficient mouse model, we demonstrate that male subfertility is caused by sterile epididymitis characterized by immune cell infiltration and extensive fibrosis. In addition, we reveal that the presence of autoreactive immune cells and inflammation in epididymides of *Aire-*deficient mice are required for iron (Fe) deposition in the interstitium, which is brought on by macrophages. We further demonstrate that male subfertility is associated with a decrease in metals zinc (Zn), copper (Cu), and selenium (Se) which serve as cofactors in several antioxidant enzymes. We also show increase in DNA damage of epididymal sperm of *Aire^−/−^* animals as a key contributing factor to subfertility. The absence of *Aire* results in autoimmune attack of the epididymis leading to fibrosis, Fe deposition, and Cu, Zn and Se imbalance, ultimately resulting in sperm DNA damage and subfertility. These results highlight the requirement of *Aire* to promote immune tolerance throughout the epididymis, disruption of which causes an imbalance of inorganic elements with resulting consequence on male fertility.

**Key points:** Breakdown of epididymal self-tolerance promotes disruption of inorganic elements. Autoimmunity causes interstitial fibrosis resulting in sperm DNA damage and subfertility. Elevated interstitial iron and macrophages contribute to fibrosis.

## Introduction

Infertility, a condition in which a couple is unable to achieve pregnancy after 12 months of unprotected intercourse, affects approximately 1 in 6 individuals worldwide, with lifetime prevalence ranging from 10.8 to 23.4% depending on global region.^1^ Male factors are thought to account about 20% of cases, and contribute to as much as 60%.^2^ Male infertility can stem from defective testicular function resulting in abnormal sperm concentration, motility, and morphology.^3^ However, many cases of infertility are unexplained, occurring despite normal sperm parameters.^4^ In these cases, underlying autoimmune disease may contribute by promoting inflammation of the reproductive organs, especially the epididymis,^5^ which could perturb sperm maturation.

Autoimmunity, or loss of immune self-tolerance, results from inappropriate T cell reactivity towards self-antigens. Self-tolerance in T cells is established in the thymus through the expression of a transcription factor called Autoimmune Regulator (AIRE) by medullary thymic epithelial cells (mTECs). In these cells, AIRE regulates transcription of tissue-restricted antigens, which are then presented in the context of major histocompatibility complex; nascent autoreactive T cells that interact too strongly with the presented self-antigen are subsequently eliminated or directed into a regulatory phenotype.^6^ In humans, mutation in the *AIRE* gene causes Autoimmune Polyglandular Syndrome Type 1 (APS-1), a rare autoimmune disease that manifests in a variety of symptoms including mucocutaneous candidiasis, hypoparathyroidism, adrenal insufficiency, gonadal insufficiency, and infertility.^7–9^

Using an Aire deficient (*Aire^−/−^*) mouse model, we previously documented CD3+ T cell infiltration into the prostate gland, seminal vesicles, epididymis, and more rarely, the testis.^10^ In addition, while sperm production and morphology appeared normal, sperm from *Aire^−/−^* males were severely impaired in their ability to fertilize eggs. *In vitro* fertilization using *Aire^−/−^*epididymal sperm resulted in only 9% of embryos reaching the 2-cell stage, and none reaching the blastocyst stage.^10^ These results suggest that malfunction of sperm in *Aire^−/−^* males are caused during post- spermatogenic maturation in the epididymis, possibly due to abnormalities in the epididymal environment.

The epididymis is an elongated, tightly coiled tube that connects each testicle to the vas deferens. In addition to storing and transporting sperm in its distal-most portion (cauda) prior to ejaculation, the epididymis provides important factors that transform the immature testicular sperm into mature, fertilization-competent sperm.^11,12^ The epididymis is indispensable for final post-testicular maturation of sperm; if they do not traverse the entire length of the epididymis, they remain incapable of fertilization.^13,14^ The rodent epididymis is divided into four anatomical regions: the initial segment (most proximal to the testis), the caput (head), the corpus (body), and the cauda (tail) (most distal to the testis, merging into the vas deferens). With their distinct transcriptomic profiles^15^ and luminal contents,^16^ each segment has specific roles in facilitating sperm maturation. The epididymis preserves sperm in a dormant state by tightly regulating the pH of the lumen.^17,18^ Epididymal epithelial cells also produce antioxidant enzymes, including glutathione peroxidases, perioxiredoxins, and superoxide dismutases, for which essential metals serve as requisite cofactors.^12,19–21 22,23^ Thus, any disruption in this epididymal environment may cause defects in sperm maturation and their fertilization competence, ultimately resulting in infertility.

Here, we demonstrate that deletion of *Aire* promotes targeting of the epididymis by autoreactive immune cells, resulting in epididymitis, interstitial fibrosis, dysregulation of inorganic physiology, and severely compromising fertility. Using a range of emerging methods to determine quantitative inorganic phenotypes,^24^ we demonstrate that autoimmune-mediated epididymal damage results in interstitial fibrosis and is associated with iron deposition, depletion of essential antioxidant cofactors such as zinc (Zn), copper (Cu), and selenium (Se), and DNA damage of the sperm. By crossing *Aire^−/−^*mice to mice deficient in T and B cells through deficiency in Recombination Activating Gene 1 (Rag1), which plays a fundamental role in generating T and B cells, we also demonstrate the role of the adaptive immune system in compromising fertility in *Aire^−/−^* males and driving fibrinogenesis and inorganic physiology.

## Materials and Methods

### Animals

Experiments with animals complied with NIH’s *Guide for the Care and Use of Laboratory Animals*,^25^ and were approved by the Institutional Animal Care and Use Committee at Michigan State University. Mice were maintained on a 12h:12h light cycle with rodent chow (Teklad 2019; Inotiv), which is formulated to support gestation in transgenic and inbred animals, and water available *ad libitum*. Mice containing mutations in the autoimmune regulator (*Aire*) gene were generated on the Balb/cJ background (Jackson Laboratories) by the Michigan State University Transgenic and Genome Editing Facility using CRISPR/Cas9 targeting. A single guide RNA (gRNA) targeting exon 2 of the *Aire* gene (**ENSMUSG00000000731**), with protospacer and PAM sequence 5’- GAC GCT CCG TCT GAA GGA GA – AGG 3’ (g158), was synthesized by IDT. Ribonucleoprotein (RNP) complexes of Cas9 and gRNA were assembled and electroporated into Balb/c zygotes to generate mutant mice as previously described.^26^ Resulting edited founders carrying 32 base-pair (bp), 44bp, or 86bp deletions were identified and backcrossed with Balb/c mice to establish the transgenic lines. Preliminary findings showed profound male subfertility in all three lines; data used in this manuscript were generated from the 32 bp line.

*Aire^+/-^* mice were crossed with *Rag1*-deficient animals to generate mice that were heterozygous for both *Aire* and *Rag1* genes. Mice were genotyped by PCR analysis of tail biopsy genomic DNA using the following primers: Aire F1: 5’-CAC ACC CTA TGC CCC CTA AC-3’, Aire R1- 5’- ACA TGG GCC GAC TTG TAT CC-3’; Rag1MutFwd: 5’-TGG ATG TGG AAT GTG TGC GAG-3’; Rag1Common: 5’-TAC ACC TTG GCG ACG ACT CCT-3’, Rag1WTFwd: 5’-TCT GGA CTT GCC TCC TCT GT-3’. *Aire*-deleted animals were confirmed for the absence of Aire protein in the thymus by immunofluorescent staining, using wild type thymus as a control.

### Fertility trial

6-week-old male *Aire*^+/+^ (n=4) and *Aire*^−/−^ (n=8) animals were paired with 6-week-old Balb/c virgin females to assess fertility. These animals were either WT or heterozygous for the *Rag1* gene. To assess the role of the adaptive immune system in fertility outcome of *Aire*^−/−^ mice, additional pairings of *Aire*^−/−^*Rag1*^−/−^ males (n=4) and *Aire^+/+^;Rag1^−/^* males (n=3) were established. Mice remained paired for 14.5 to 16 weeks, with the exception that two of the three *Aire^+/+^;Rag1^−/−^* males were paired for ∼6 weeks. At the conclusion of the fertility trial, males were euthanized for downstream analyses at 19 to 23 weeks of age; additional males were euthanized at 7-9 weeks. Females were euthanized at the same time as the males, except when they were pregnant, in which case they were euthanized following birth of the litter. Fertility rate was calculated as:

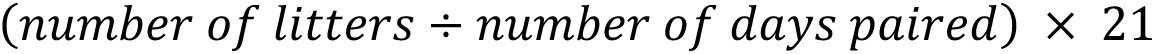

We chose 21 days as an expected length of mouse gestation; if dams produced a litter of pups every 21 days, the resultant number would be close to 1.

### Histology

Paraffin-embedded tissues were fixed in 4% paraformaldehyde at 4°C overnight, transferred to 70% ethanol, embedded in paraffin, and sectioned at 5 µm onto SuperFrost slides (Thermofisher Scientific). Alternatively, tissues were fixed in 4% paraformaldehyde at 4°C overnight, transferred to 30% sucrose, and embedded in OCT medium (Sakura) by freezing the medium in liquid-nitrogen chilled isopentane. OCT-embedded samples were kept at -80°C until cryosectioning (Leica Biosystems). For hematoxylin and eosin staining,^27^ we used the Leica SelecTech staining system consisting of Hematoxylin 560, Blue Buffer 8, Define, Alcoholic Eosin Y 515. Sections were deparaffinized in Histoclear (National Diagnostics) and rehydrated in a gradient of ethanol (100%, 95% and 80%) and water. Slides were stained in hematoxylin for 3 minutes, washed in water for 1 minute, and immersed in 95% ethanol for 40 seconds followed by alcoholic eosin Y for 12 seconds. Slides were then washed using two exchanges of 90% ethanol for 40 seconds and dehydrated in 100% ethanol for 5 minutes. Finally, slides were washed in Histoclear for 5 minutes and mounted using Cytoseal 60 (Fisher Scientific). For Masson’s trichrome staining, which was used to assess fibrosis, serially sectioned samples were submitted to the Investigative Histopathology Core at Michigan State University. Using this stain, connective tissue stains blue, nuclei stain purple/red, and cytoplasm stains pink/red.

### Immunohistochemistry and Immunofluorescence

Paraffin-embedded epididymal tissue sections were immunohistochemically stained with rabbit anti-CD8 (ab217344, Abcam), rabbit anti-CD4 (Abcam), rat anti-CD19 (6OMP31, Thermofisher Scientific), rat anti-F4/80 (123102, Biolegend), or rabbit anti-αSMA (14395-1-AP, Proteintech) antibodies. After deparaffinization and rehydration, we performed a heat-mediated antigen retrieval step using Reveal Decloaker sodium citrate buffer (Biocare Medical), heated to 199°F for 20 minutes. After cooling to room temperature, samples were incubated with the SuperBlock™ blocking buffer (37515, ThermoFisher Scientific) for 10 minutes followed by 3% hydrogen peroxide in methanol to quench endogenous peroxidase activity for 10 minutes. Sections were washed with phosphate buffered saline (PBS) before incubating with the primary antibody. Adjacent sections were used for isotype controls. Biotinylated anti-rabbit or anti-rat IgG secondary antibody (Vector Laboratories) was applied, followed by Ready-To-Use Horseradish Peroxidase and DAB (Vector Laboratories) for color development.

To quantify positively stained cells, slides were scanned using a Zeiss Axioscan 7 Microscope Slide Scanner (Carl Zeiss AG). Three nonoverlapping, randomly chosen regions of interest (ROI; 350 µm ξ 950 µm) were from the caput, corpus, and cauda epididymides were captured using Zen Microscopy Software (Carl Zeiss). In some animals, only two non-overlapping regions could be imaged. Three independent blinded evaluators counted DAB+ cells from slides stained with anti- CD4 and anti-CD8 antibodies. The non-parametric Wilcoxon’s t-test was used to compare the means of DAB+ cells between *Aire^+/+^* and *Aire^−/−^* samples. Intraclass Correlation Coefficient was used to assess the strength of inter-evaluator agreement and was calculated using the “icc” R package. The ICC for this experiment was 0.91, indicating a high similarity between values measured by the three independent evaluators.

For immunofluorescence, OCT-embedded samples were sectioned at 5 μm, washed in PBS for 10 minutes, blocked using SuperBlock Blocking Buffer, and incubated overnight with antibodies against F4/80, CD163 (1:400; Abcam), and/or Ferritin (1:1000; F5012, Sigma Aldrich). The following day, the slides were washed and incubated with secondary antibodies (AF488-anti-rat or AF546-anti-rabbit IgG; 0.01mg/mL, ThermoFisher Scientific) and mounted using ProLong Gold antifade reagent with DAPI (ThermoFisher Scientific).

### Flow cytometry

For flow cytometeric analysis of lymphocytes, epididymides from 2 males were pooled into 1 sample, such that 1 biological replicate consisted of cells from 2 males and 4 epididymides. Once excised, cauda epididymides were dissected from the rest of the epididymis and collected into TYH media warmed to 37°C to allow sperm to swim out of the sample. Cauda epididymides were further processed and analyzed separately from caput-corpus epididymides such that spatial information could be maintained while ensuring sufficient cellular yield. Tissues were finely minced using spring scissors, collected into gentleMACS™ M Tubes (Miltenyi Biotec), and incubated in 5 mL of Miltenyi Tissue Dissociation Kit 1 (Miltenyi Biotec) for 15 minutes at 37°C, using GentleMACS™ Dissociator (Miltenyi Biotec) for 1 minute per sample at room temperature. The samples were returned to 37°C to incubate for a 15 additional minutes. The digest was filtered through a 70 µm filter into 50 mL conical tubes containing 10 mL of PBS with 5% FBS to neutralize the digestive enzyme. Each sample was topped up to 30 mL with 5% FBS and centrifuged at 400g for 5 minutes. The cell pellet was washed with 10 mL of PBS/5%FBS buffer, resuspended in 0.4 mL of PBS/5%FBS buffer, and aliquoted into 96 well plates for staining. All cells were incubated with a viability stain (1:1000; Invitrogen™ Live/Dead™ Fixable Blue Dead Cell Stain kit, ThermoFisher Scientific) for 30 minutes on ice, followed by staining with Mouse BD FcBlock™ (1:100, 553142BD Pharmingen, Cat. No.) for 20 minutes on ice. Cells were pelleted and resuspended with the primary antibody cocktail on ice for 30 minutes. The following antibodies were used: BV421 anti-CD45 (1:200; 103133, 30-F11, Biolegend,), KIRAVIA Blue 520 anti-CD4 (1:100, clone GK1.5; Biolegend), BV785 anti-CD8 (1:100, 53-6.7, Biolegend), PE-Dazzle 594 anti-CD44 (1:100, IM7, Biolegend), PE-Cy5 anti-CD69 (1:50; H1.2F3, Biolegend), APC anti-CD25 (1:50; PC61, Biolegend), BV510 anti-B220 (1:100; RA3-6B2, Biolegend). After incubation with primary antibodies, the cells were washed three times in 200 µL of flow staining buffer, pelleting the cells between each wash. Lastly, cells were fixed using 1% paraformaldehyde in PBS for 15 minutes on ice and stored at 4°C until analysis using Cytek® Aurora spectral flow cytometer (Cytek® Biosciences,).

### Gelatin standard preparation

Gelatin standards (blank, 1, 5, 10, 20, 50 parts per million (ppm)) (**Supplemental Table 1**) were produced by mixing in IV-Stock-4 (Inorganic Ventures) to 10% porcine gelatin (G2500-500G, Sigma Aldrich) and heated to 55°C. The blank contained no IV-Stock-4. 400 µL of the preparation was pipetted directly onto the cryostat mounting chuck, left to freeze at -20°C for 4 minutes, and sectioned at 20 μm thickness onto the SuperFrost glass slides (Thermofisher Scientific). A 50 µL aliquot of each standard was aliquoted in a pre-weighed metal-free tube to determine elemental concentrations using an Agilent 8900 Triple Quadrupole - Inductively Coupled Plasma – Mass Spectrometer (ICP-QQQ-MS, Agilent) equipped with the Agilent SPS 4 Autosampler, integrate sample introduction system (ISiS), x-lens, and micromist nebulizer. To each sample, 300 µL of trace metal grade 70% nitric acid (Fisher chemical, #A509P212), was added to digest the sample at 70 °C for 4 hours. Following digestion, 9.7 mL of ultrapure deionized water was added to each sample. All samples were weighed prior to analysis on the ICP-QQQ-MS.

### Spatial, quantitative mapping element distribution using LA-ICP-TOF-MS

Paraffin-embedded 5 µm tissue sections were mounted onto charged glass slides and laser ablated using the Elemental Scientific Lasers BioImage 266 nm laser ablation system (Elemental Scientific Lasers), which is equipped with an ultra-fast low dispersion TwoVol3 ablation chamber and a dual concentric injector (DCI3). The aerosolized sample was transferred to the Tofwerk icpTOF S2 mass spectrometer (TOFWERK AG) where the elemental content was analyzed real time according to mass/charge (m/z) ratio. Daily tuning was performed using NIST SRM612 glass certified reference material (National Institute for Standards and Technology). High intensities for ^140^Ce and ^55^Mn were used to optimize torch alignment, lens voltages, and nebulizer gas flow while maintaining low oxide formation based on the ^232^Th^16^O+/^232^Th+ ratio (< 0.5). Parameters are listed in **Supplemental Table 2**. Laser scanning was performed in a raster pattern with a laser repetition rate of 100 Hz using a 10 μm circular spot size with laser fluence set to 15 J/cm^2^. Ten lines of gelatin standards were ablated using the same parameters but with the raster spacing set at 20 μm. Scanning data was recorded using TofPilot 1.3.4.0 (TOFWERK AG) and saved in the open-source hierarchical data format (HDF5). Post-acquisition data processing was performed with Tofware v3.2.0, a data analysis package used as an add-on to IgorPro (Wavemetric Inc.). Imaging and mass calibration was conducted using the Iolite version 4.8.6. Software (Elemental Scientific Lasers) .^28^

### Elemental Composition Determination by ICP-QQQ-MS

Mouse epididymides separated into caput, corpus, and cauda were placed into a pre-weighed metal-free 15mL conical tubes (Labcon) to be analyzed using ICP-QQQ-MS (Agilent). Prior to analysis, each metal-free tube was weighed using an analytical balance pre- and post-addition of the tissue sample to ascertain the weight of the tissue. Tissues were digested in 300 µL of 70% nitric acid (Fisher, #A509P212) and incubated at 65°C overnight. The following day, 9.7 mL of ultrapure deionized water was added to bring the final nitric acid concentration to 3% by volume. Each tube was weighed again before analysis to ascertain the final weight of the sample. Standards were prepared using IV-Stock-4 (Inorganic Ventures) at 0.01, 0.1, 1, 10, and 100 parts per billion (ppb) using 3% nitric acid in metal-free 15 mL conical tubes. The element concentrations of the samples were calculated using the following equation:

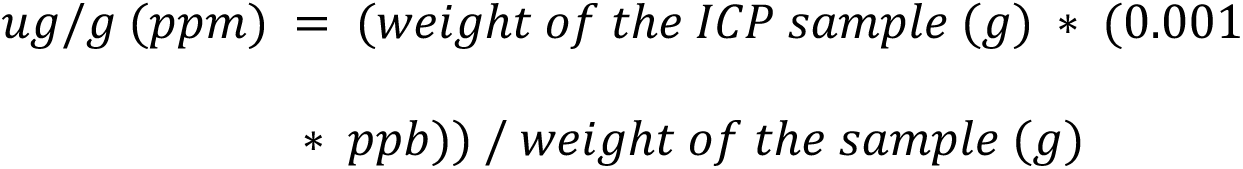

### Comet assay

The assay was conducted using the CometAssay® Single Cell Gel Electrophoresis Assay kit (Catalog # 4250-050-K, R&D Systems). Cauda epididymides were collected from *Aire^+/+^* and *Aire^−/−^* males into a 1.5 mL tube filled with 250 µL of PBS, gently minced with spring scissors, and incubated for 15 minutes at 37°C to allow the sperm to swim out of the tissue. The supernatant was collected into a separate 1.5mL centrifuge tube and counted using a hemocytometer by mixing 5 µL of the suspension with 95 µL of MilliQ water. Fifty µL of the resuspended sperm (1 million cells/mL) were mixed with 500 µL of 1% LMAgarose, spread onto a glass slide, and placed at 4°C for 30 minutes. The samples were then incubated with lysis buffer + Proteinase K (500 µg/mL) for 18 hours at 4°C. The following day, the slides were immersed in cold Neutral Electrophoresis Buffer (0.5M Tris-HCL, 1.5M sodium acetate) and subjected to electrophoresis at 21V/cm for 45 minutes at 4°C. Next, the slides were immersed in DNA Precipitation Solution and immersed in 70% ethanol for 30 minutes each at room temperature. The slides were dried at 37°C and stained with 100 uL of SYBR® Gold Nucleic Acid Gel stain (1:10000, S11494, ThermoFisher Scientific) for 30 minutes at room temperature. Lastly, the slides were briefly washed in tap water, dried, and imaged using a 488 nm filter (Zeiss Axioscan 7). To calculate the percentage of sperm with comet tails, Fiji software was used to convert the images to 8-bit, and threshold was set between 65-70 range to 255, followed by the “Analyze Particle” function in Fiji, which counted the number of sperm heads (particle size set to 10 - 250) present on the image. To calculate proportion of sperm with fragmented DNA, the comet tails were manually counted from the same image, divided by the total number of particles detected, and multiplied by 100.

### Statistical analysis

R studio and the packages ggplot2, ggbeeswarm, readxl, ggpubr, ggsci, icc, and tidyverse were used to generate graphs and conduct statistical analysis. Normal distribution of the data was tested using the Shapiro-Wilks test. For comparison of the means between two independent, normally distributed samples, Student’s t-test was used; for more than two, one-way ANOVA followed by Tukey’s post-hoc test was used. For samples that failed Shapiro-Wilks normality test, Wilcoxon’s test was used to compare the means of two samples, and Kruskal-Wallis test for two or more samples, followed by Dunn’s post-hoc test. For the comet assay, a paired t-test was conducted. *P*- values less than 0.05 were deemed statistically significant.

## Results

### *Aire^−/−^* males subfertility is dependent on the adaptive immune system

We generated transgenic animals heterozygous for *Aire* and *Rag1* genes on the Balb/c background. The absence of *Rag1* gene with *Aire*-deficiency (*Aire^−/−^*) produces animals without adaptive T and B cells (*Aire^−/−^;Rag1^−/−^*), allowing us to address the role of central tolerance to male reproductive tissues. Since we rederived the *Aire^−/−^* animals using CRISPR/Cas9, we validated the absence of Aire protein in the thymus using immunofluorescence **(Fig 1A)**. We also tested whether the *Aire*^−/−^ animals recapitulated the development of autoimmunity and infertility, as previously demonstrated by us and others.^6,10,29^ We observed immune cell infiltration in both reproductive and non-reproductive organs (**Fig. 1B**), confirming that our *Aire*^−/−^ mouse model recapitulated established mouse models of APS-1.^6^ Because we observed smaller testes with immune cell infiltration in about 14% of *Aire*^−/−^ males examined,^10^ we euthanized 10- and 20-week-old *Aire*^−/−^ males to assess testicular histology. Hematoxylin and eosin staining showed normal seminiferous tubules with intact spermatogenesis, comparable to *Aire*^+/+^ testes in a majority of our *Aire*^−/−^ samples at 10 weeks of age (*Aire^+/+^*; n=4, *Aire^−/−^*; n=4) and 20 weeks of age (*Aire^+/+^*; n=9, *Aire^−/−^*; n=10, data not shown), suggesting that testes were not targeted by immune cells, or that the testes- blood-barrier prevented immune cell infiltration in these *Aire^−/−^* mice.

**Figure 1.**
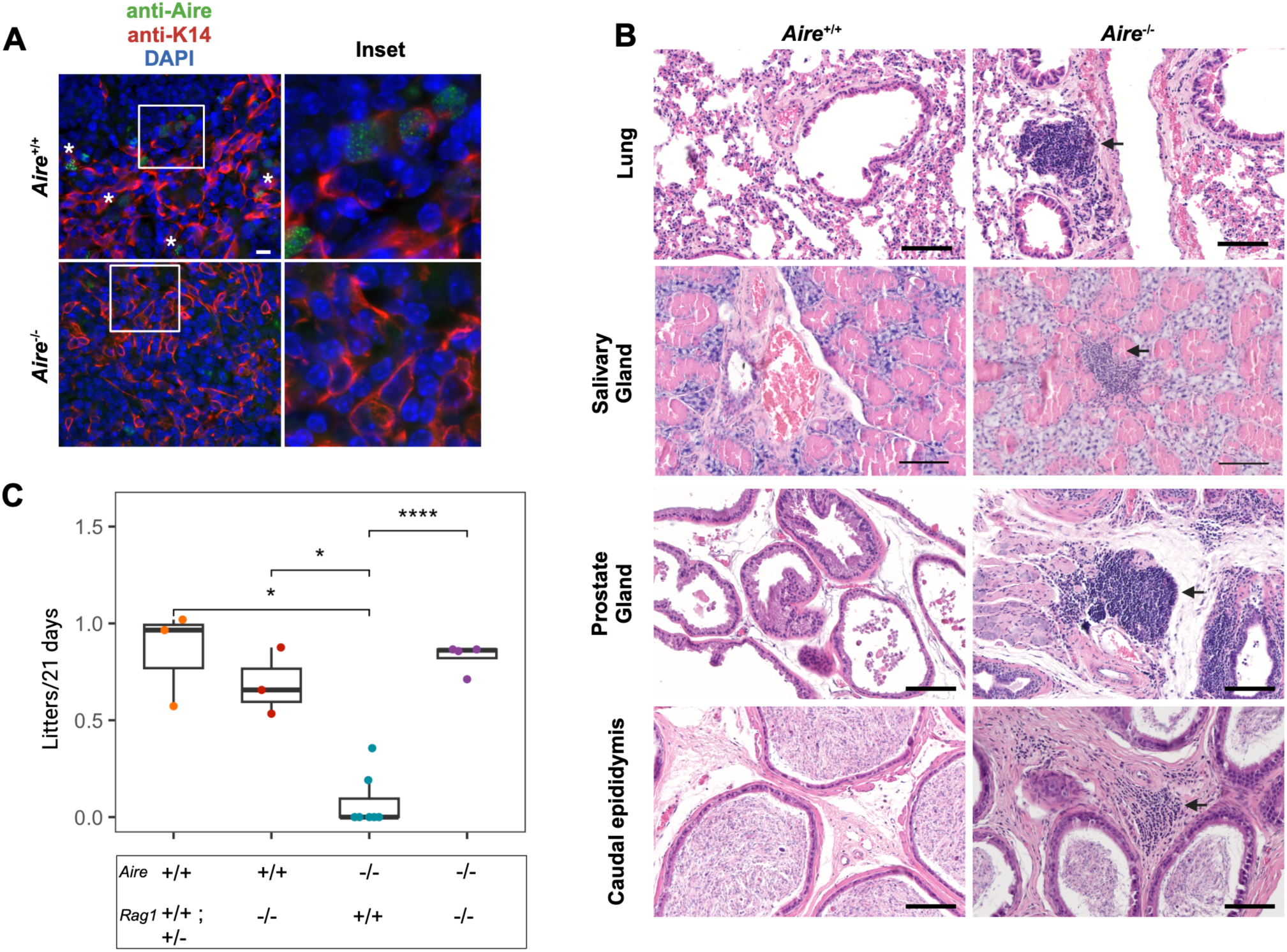
Aire-deficient (*Aire*^−/−^) males develop autoimmunity towards non-reproductive and reproductive organs and are subfertile. (A) Immunofluorescence of Aire in *Aire^−/^*^−^ and *Aire*^+/+^ thymi. AF488 (green), Aire; AF568 (red), Cytokeratin 14 (K14, mTEC marker); DAPI (blue), nuclei. Asterisks highlight areas in which additional Aire^+^ cells can be seen outside the inset. Note the absence of the characteristic intranuclear speckles in the *Aire^−/−^* thymus. Scale bar = 10μm. (B) Representative hematoxylin and eosin-stained lung, salivary gland, prostate, and cauda epididymis sections from 20 week-old *Aire^−/^*^−^ and *Aire^+/^*^+^ males. Lung: n=3 per genotype; prostate: *Aire^−/−^,* n=11/*Aire^+/^*^+^, n=7; caudal epididymis: n=11 per genotype. Arrows point to lymphoid aggregates. Scale bars = 100μm. (C) Fertility outcomes of *Aire*^−/−^ and *Aire^+/+^* males on the *Rag1*-sufficient or - deficient background. ANOVA followed by Tukey’s post hoc test. ns: *P*>0.05, \**PP* < 0.0001.

To test fertility, 6-week-old *Aire*^+/+^ (n=3) or *Aire*^−/−^ (n=7), which were either heterozygous or wild-type for the *Rag1* gene, were singly housed with virgin WT Balb/c females. Additionally, fertility of *Aire^+/+^*;*Rag1^−/−^* (n=3) and *Aire^−/−^;Rag1^−/−^*(n=4) males was evaluated. Wild type (*Aire*^+/+^*Rag1^+^*) males demonstrated robust fertility, with an average delivery rate of 0.85 litters per 21 days and 74 pups total over ∼14 weeks. In contrast, the *Aire^−^*^/-^*;Rag1^+^*males had an average delivery rate of 0.08 litters per 21 days, with only 17 total pups produced. The two litters produced by the seven *Aire^−/−^* males occurred in the first 7 weeks of pairing, whereas *Aire^+/+^* males were able to sire litters for the entire duration of pairing. *Aire^−/−^;Rag1^−/−^* males showed fertility comparable to *Aire^+/+^* mice **(Fig. 1C),** showing that adaptive immune cells are responsible for subfertility in *Aire*^−/−^

### *Aire*^−/−^ epididymides show immune cell infiltration

Because the testis of *Aire^−/−^* mice appeared normal, we focused our attention on the epididymis, where the testicular sperm undergo maturation, gain fertilization competency, and are stored before ejaculation. Histological analysis of 10 and 20-week-old *Aire*^−/−^ epididymides revealed thickening of the interstitium and immune cell infiltration consisting of CD4+ and CD8+ T cells, and CD19+ B cells **(Fig. 1B, 2A).** To measure the degree of infiltration of T cells, we counted the immunohistochemically stained cells, with attention to age-dependent variation and differences between the epididymal caput (head), corpus (body), and cauda (tail) (**Fig. 2B**). While CD4 and CD8 cells in *Aire^+/+^* epididymides were found only sporadically, both cell types were frequently found in *Aire^−/−^*mice in all three segments of *Aire^−/−^* animals in both young and old animals. No differences between regions of epididymis within each strain were observed (*P*>0.05). Age-dependent differences were not obvious, therefore data from young and aged animals were combined. T cell infiltration of all three epididymal regions of *Aire^−/−^* mice were significantly higher than those of *Aire^+/+^* mice (**Fig. 2C, D**). As expected, neither T nor B cells could be identified in *Rag1*-deficient mice, regardless of absence or presence of *Aire* (not shown).

**Figure 2.**
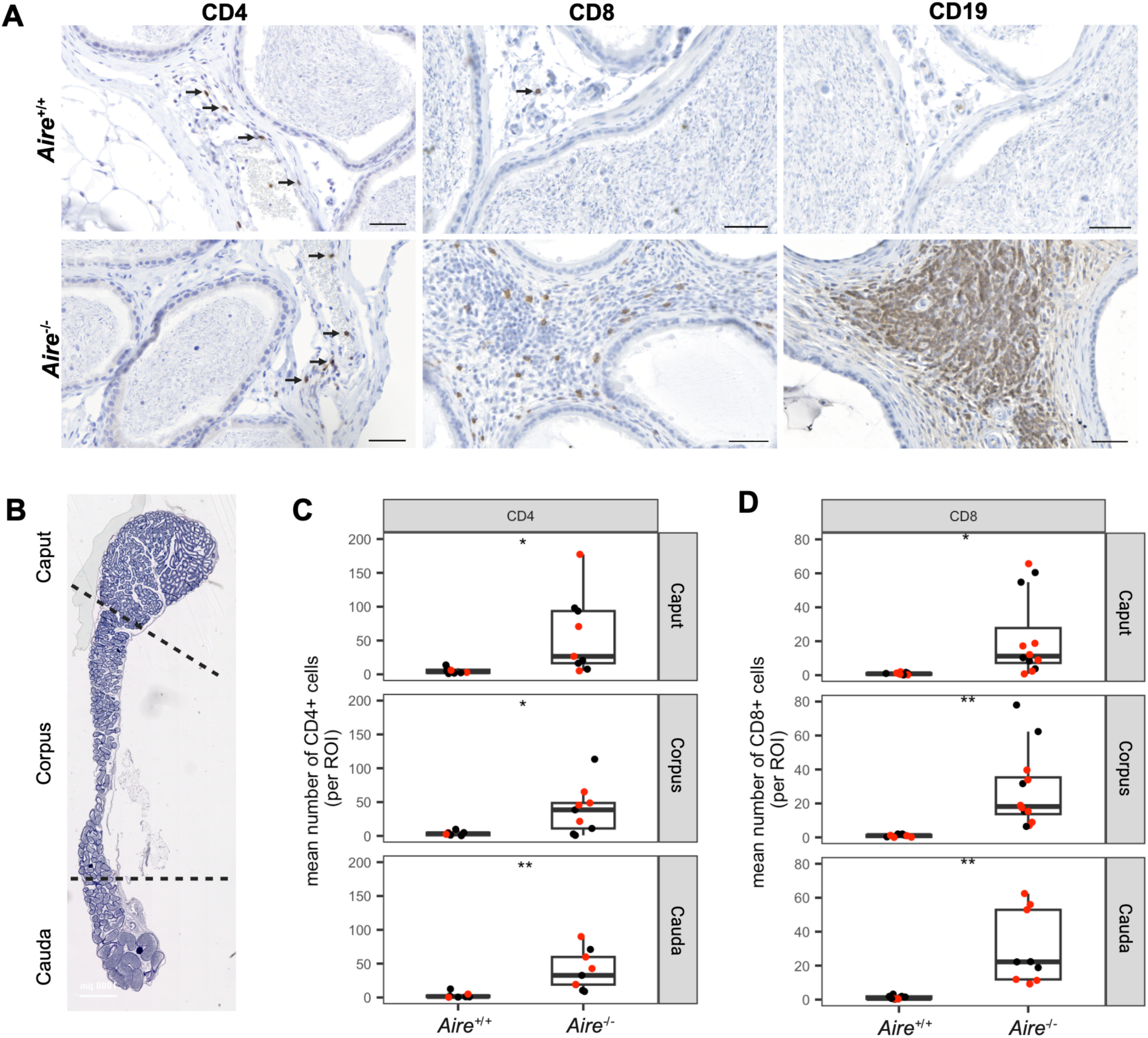
Immune cell infiltrates consisting of CD4+ T cells, CD8+ T cells and CD19+ B cells occur early and persist with age in *Aire^−/−^* epididymides. (A) Immunohistochemical staining for CD4+ T cells, CD8+ T cells and CD19+ B cells in the epididymis of 19-20 week old *Aire^+/+^* and A*Aire^−/−^* epididymides. CD4+ and CD8+ T cells are also found sporadically in the *Aire^+/+^* epididymis (arrows). *Aire^+/+^*(19-20 weeks: n=5), and *Aire^−/−^* (19-20 weeks: n=7), Scale bar = 50 μm. (B) Photomicrograph of the three main anatomical regions of the epididymis (Caput, H; Corpus; B; Cauda; T). (C, D) Manually enumerated CD4+ (*Aire^+/+^*: n=8; *Aire^−/−^*: n=9) and CD8+ (*Aire^+/+^*: n=10, *Aire^−/-:^* n=12) T cells in the epididymis. Red dots denote 19-20 week old animals; black dots denote 7-9 week old animals. Statistical Analysis: Wilcoxon non-parametric t-test. ns: p>0.05, \**PP*<0.01.

To characterize the activation status of the infiltrating epididymal T cells, we performed flow cytometry, separating the caput and corpus from the cauda epididymides. Within genotype, no differences were observed in the relative abundance of CD45hi immune cells in the different regions of the epididymis (*P*>0.05). Between genotypes, the proportion of CD45hi cells among total cells trended towards increasing in the corpus-caput, and significantly increased in the cauda epididymis, of *Aire^−/−^* mice as compared to *Aire^+/+^* controls (**Fig. 3B**). Similarly, the proportion of B cells remained similar within genotype, but increased in the cauda epididymis of *Aire^−/−^* mice (**Fig. 3C**). Relative abundance of CD4+ and CD8+ T cells was higher in the epididymal caput- corpus of *Aire^−/−^* mice than that of *Aire^+/+^*controls (**Fig. 3D, E**). In the cauda epididymis, CD4+ T cells remained similar between knockout and control mice (**Fig. 3F**), but CD8+T cells increased approximately 6-fold in *Aire^−/−^*mice **(Fig. 3G)**. Finally, the early activation marker CD69 was elevated on CD8+ T cells of both caput-corpus (**Fig. 3E**) and cauda (**Fig. 3G**); in CD4+ T cells, the CD69 expression appeared to increase but this did not reach statistical significance, possibly due to insufficient statistical power (**Fig. 3D, F**). Nearly all CD4+ and CD8+ T cells were CD44^high^, indicating prior antigen experience; levels of CD44, however, were unchanged between genotypes (**Fig. 3D-G**). CD25 expression was also unchanged bewteen *Aire^−/−^*and *Aire^+/+^* mice, except for CD4+ cells obtained from cauda epididymis (**Fig. 3F**). Few CD8+ T cells in either genotype expressed IL-2Rα/activation marker CD25 (**Fig3. E, G**).

**Figure 3.**
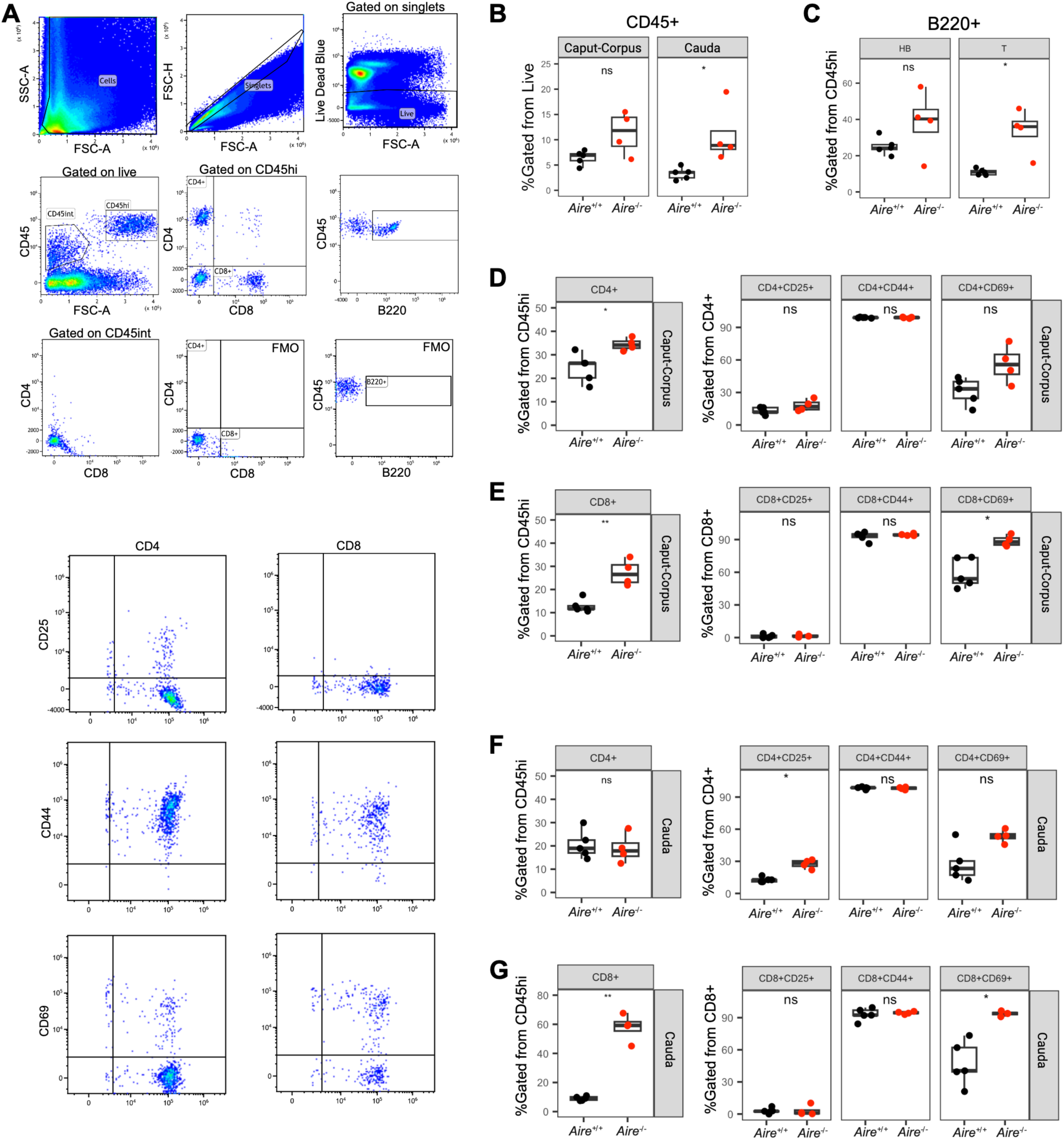
Flow cytometric analysis of CD4+ and CD8+ T cells from *Aire^−/−^* and *Aire^+/+^*epididymides. (A) Gating strategy for flow cytometric assessment of lymphocyte population in the epididymis. FSC-A: forward scatter area, SSC-A: side scatter area, FSC-H: forward scatter height. Epididymides were separated into Caput-Corpus and Cauda, digested into single cell suspensions, and analyzed by flow cytometry for the expression of CD25, CD44, and CD69. (B) Relative abundance of CD45hi cells in *Aire^+/+^* and *Aire^−/−^*epididymides; (C) Relative abundance of CD45hi B220+ cells in the epididymis; (D, F) Relative abundance of CD45hi CD4+T, followed by CD4+CD25+ CD4+CD44+, and CD4+CD69+ T cells in the epididymis; (E,G) Relative abundance of CD45hi CD8+ T cells, followed by CD8+CD25+, CD8+CD44+, and CD8+CD69+ T cells in the epididymis. Animals analyzed were 19-20 weeks old. Wilcoxon t-test, ns: *P*>0.05, \**P*< 0.01.

### 56Fe is increased in *Aire^−/−^* epididymis and correlates with fibrosis

Thickening of the epididymal interstitium in *Aire^−/−^*mice (**Fig. 1B**) could indicate fibrotic change of the tissue. To test this, tissues were stained with Masson’s trichrome, which revealed increased presence of collagen fibers in *Aire*^−/−^ epididymides **(Fig. 4)**. We also probed for αSMA+ myofibroblast using immunohistochemistry, which are pertinent to fibrogenesis and scar formation^30^ (**Supplemental Fig. 1**). These cells were abundant in the interstitium of *Aire^−/−^*epididymis compared to *Aire^+/+^*, indicating their potential involvement in fibrogenic transformation of the interstitium in our *Aire^−/−^* animals. The epididymis of the age-matched *Aire^−/−^;Rag1^−/−^*animals, which possess innate immune cells but lack mature T and B lymphocytes, did not develop fibrosis (**Fig. 4, third column)**, demonstrating that autoreactive adaptive immune cells are required to promote fibrogenesis of the epididymis.

**Figure 4.**
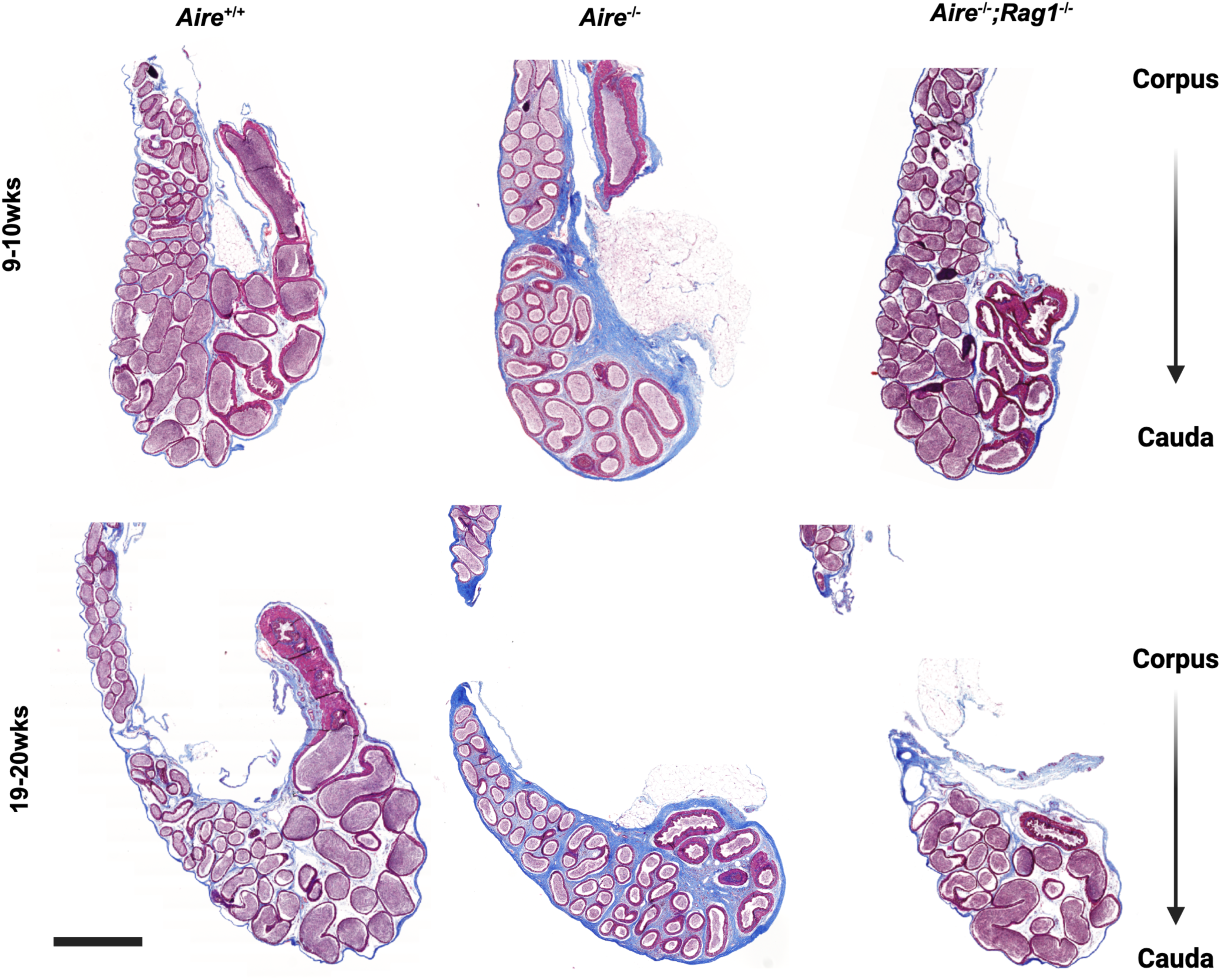
Fibrotic changes in epididymides of *Aire^−/−^* males. Masson’s trichrome staining reveals extensive fibrosis (blue staining) in the interstitium of *Aire^−/−^* epididymides of 9-10 weeks and 19- 20-week-old males compared to that of *Aire^+/+^*and *Aire^−/−^;Rag1^−/−^* controls. Shown are the cauda and portions of the corpus epididymis. Scale bar = 1000μm.

Dysfunction in iron metabolism and homeostasis can cause hemochromatosis and idiopathic pulmonary fibrosis, both of which are characterized by extensive fibrotic changes of the organ.^31,32^ To determine whether iron accumulation contributed to fibrosis in *Aire^−/−^* epididymis, we dissected the epididymis into its anatomically distinct regions **(Fig. 2B)** and used ICP-QQQ-MS for elemental quantification. Using this method, we found that ^56^Fe concentration (μg/g tissue) regardless of age was elevated throughout the epididymis from *Aire^−/−^* males compared to *Aire^+/+^*and *Aire^−/−^;Rag1^−/−^* **(Fig. 5A)**. In *Aire^+/+^* epididymides, the mean concentration of ^56^Fe was comparable between caput, corpus, and cauda (21.5 ± 5.2 µg/g; 26.8 ± 10.1 µg/g; 29.8 ± 4.9µg/g, respectively). However, in *Aire^−/−^* epididymides, concentrations were 3-4-fold higher (caput, 69.8 ± 51.3 µg/g; corpus, 110.1 ± 35.7 µg/g; cauda, 85.2 μg/g ± 16.0 µg/g). The ^56^Fe concentration in each of the epididymal segments of the *Aire^−/−^;Rag1^−/−^*tissues were comparable to *Aire^+/+^* tissues and values were not significantly different (**Fig. 5A)**. ^56^Fe concentration in the caudal sperm fraction was similar between *Aire^+/+^*and *Aire^−/−^*, suggesting that increased ^56^Fe is localized to the tissue rather than the sperm **(Fig. 5B).**

**Figure 5.**
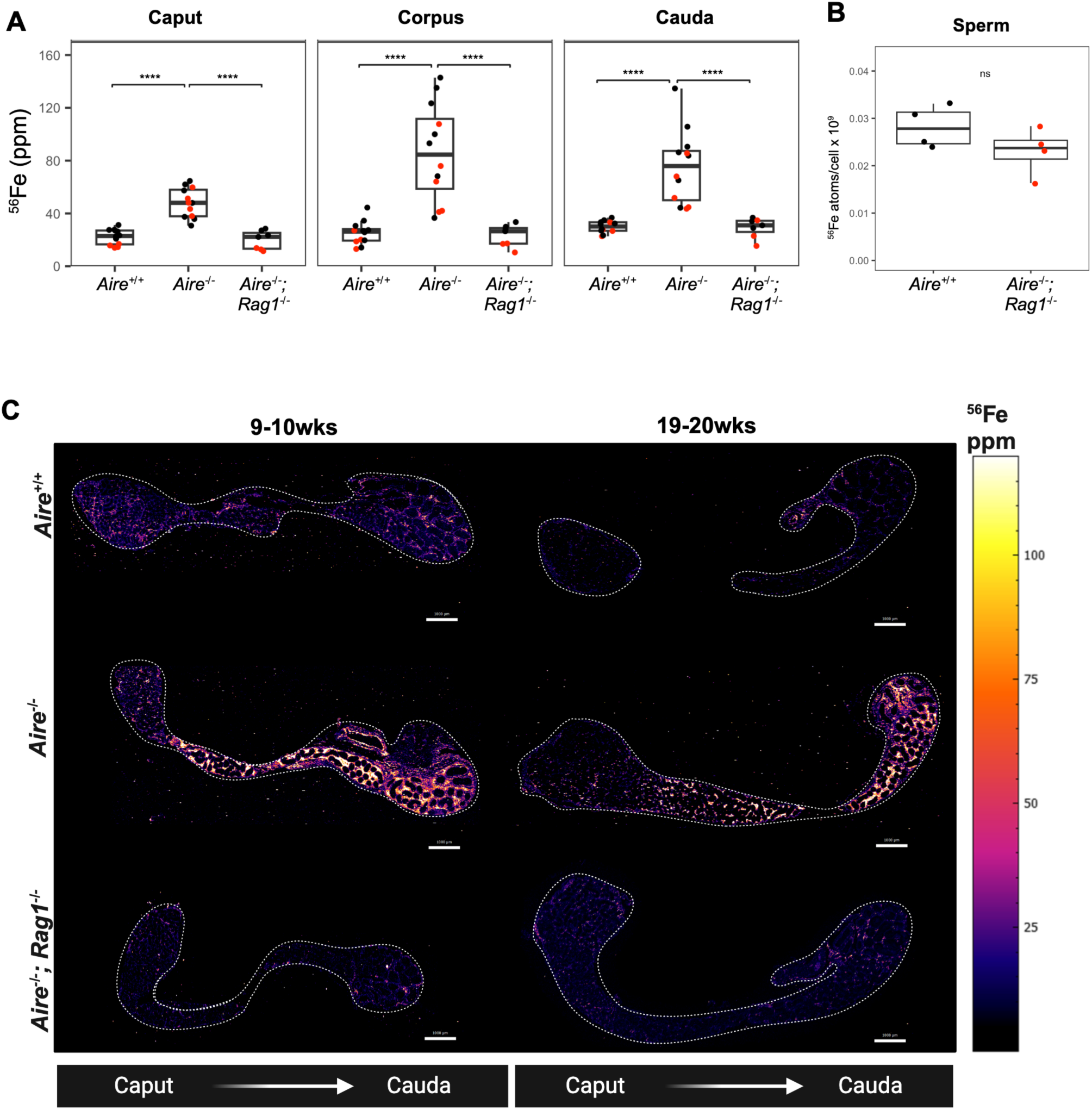
^56^Fe concentration is increased in the *Aire^−/−^* epididymis and localizes to the interstitium. **(A)** ICP-QQQ-MS analysis of epididymal segments. Red = 7-9 wks, Black = 19-20 wks. **(B)** ^56^Fe measurement in sperm using ICP-QQQ-MS. **(C)** LA-ICP-TOF-MS reveals the spatial distribution of ^56^Fe to the interstitium of the *Aire^+/+^*(n=3 for each age), *Aire^−/−^* (n=4 for each age), and *Aire^−/−^;Rag1^−/−^*(n=4 for 9-10wks, n=3 for 19-20wks) epididymis. Statistical analysis: Kruskal-Wallis One-Way ANOVA followed by Dunn-Bonferroni post-hoc was conducted to compare the means of each group for (A). Wilcoxon-test for (B). \*\*\*\**P*<0.00005, ns = not significant.

To visualize the spatial distribution of iron in the epididymis, we used laser ablation – inductively coupled plasma – time of flight – mass spectrometry (LA-ICP-TOF-MS) **(Fig. 5C**). Analogous to the spatial single-cell RNA sequencing that illuminates transcriptomics of a sample on a per-pixel basis, LA-ICP-TOF-MS can delineate the total elemental content of a sample and instantaneously allow spatial visualization of all elements within the periodic table in the tissue sample. Using this technique, we found that ^56^Fe was especially concentrated within the interstitium of the corpus and cauda regions of both the young and the aged *Aire^−/−^* epididymides; this was not observed in either *Aire^+/+^* or *Aire^−/−^;Rag1^−/−^* tissues **(Fig. 5C).**

### F4/80+ macrophages are sources of ^56^Fe in the interstitium

We next asked what cells the excess iron originates from in *Aire^−/−^* epididymides. Because of known associations between macrophages and iron in fibrosis of the lung,^33^ we hypothesized that these cells are a source of iron. We first used a macrophage-specific marker, F4/80, to characterize the distribution of macrophages in the epididymis of *Aire^+/+^*, *Aire^−/−^*and *Aire^−/−^;Rag1^−/−^* animals using immunohistochemistry (**Fig. 6A**). In the control tissues, F4/80+ macrophages were found as small, densely stained cells in the interstitium of the epididymis. In *Aire^−/−^*tissues, macrophages became elongated and were found throughout the interstitium of *Aire^−/−^* epididymis (**Fig. 6A),** corresponding with the locations of fibrosis and iron accumulation (**Fig. 4**, **Fig. 5**). F4/80+ macrophages found within fibrotic regions of the epididymis were also CD163+ (**Fig. 6B**), a marker commonly used to identify alternatively activated macrophages with profibrotic transcriptomic signatures.^34^ Collectively, the results suggest that macrophages with a pro-fibrotic phenotype contribute to interstitial fibrosis in the *Aire^−/−^* epididymis.

**Figure 6.**
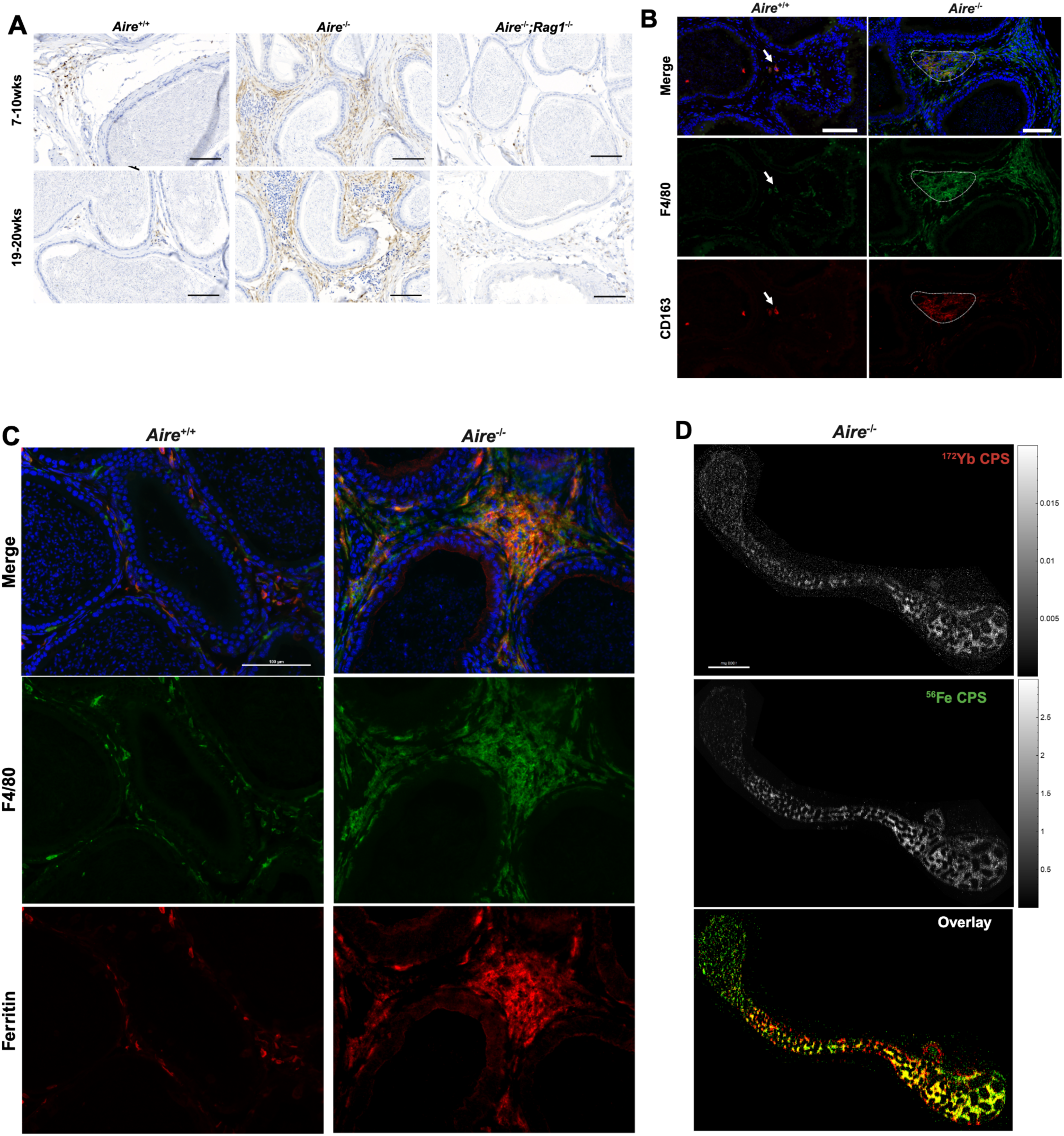
F4/80+ macrophages carry intracellular ferritin-associated iron in the interstitium of *Aire^−/−^* epididymis. **(A)** Representative immunohistochemical staining for F4/80+ macrophages in cauda epididymides of *Aire^−/−^* compared to *Aire^+/+^* and *Aire^−/−^;Rag1^−/−^ mice. Aire^+/+^*(7-10 weeks: n=4, 19-20 weeks: n=5), *Aire^−/−^* (7-10 weeks: n= 4, 19-20 weeks: n=7), and *Aire^−/−^;Rag1^−/−^* (7-10 weeks: n=4, 19-20 weeks: n=3) Scale bar = 100 μm. **(B)** Immunofluorescence staining for F4/80+ (Alexa Fluor 488, green) and CD163+ (Alexa Fluor 546, red) macrophages in *Aire^+/+^*and *Aire^−/^*^−^ epididymal tail. Scale bar = 50 μm. **(C)** Dual immunofluorescence staining for F4/80 (Alexa Fluor 488, green) and ferritin (Alexa Fluor 546, red) in *Aire^+/+^* and *Aire^−/^*^−^ cauda epididymis. Scale bar = 100 μm. (**(D)** Representative LA-ICP-TOF-MS images of *Aire^−/−^* epididymis stained with F4/80 primary antibody followed by a secondary antibody tagged to a lanthanide (^172^Yb) revealing colocalization of ^56^Fe with F4/80+ macrophages. Scale bar = 1000μm.

Due to the ability of macrophages to store iron in the form of ferritin and hemosiderin, we asked whether iron co-localized with F4/80+ macrophages. Indeed, we found that ferritin, a 24-mer iron storage protein,^35^ often co-localized with F4/80-positive macrophages, particularly in Aire^−/−^ animals (**Fig. 6C**). Interestingly, in wild type animals, ferritin was found in and F4/80 overlapped less faithfully, suggesting heterogeneity of iron regulatory mechanisms in F4/80+ macrophages, an/or the presence of iron-carrying non-macrophages such as eosinophils or neutrophils.^36,37^ To further confirm the asociation between iron and macrophages in Aire^−/−^ mice, we labelled epididymides with a primary antibody for F4/80 followed by a secondary ^172^Yb-tagged antibody, and in the same sections, produced elemental maps using LA-ICP-TOF-MS. The resulting images were co-registered on a per-pixel bases and showed co-localization of ^172^Yb-positive pixels with the ^56^Fe signature, strongly suggesting that macrophages are the source of excess intracellular iron (**Fig. 6D**).

Excess iron in tissues could lead to reactive oxygen species through the Fenton reaction,^38–40^ and in turn could cause DNA damage. Therefore, we measured DNA fragmentation in sperm isolated from caudal epididymis of *Aire^+/+^* or *Aire^+/-^* and *Aire^−/−^* males using the comet assay, which reveals single- and double-stranded breaks in the DNA. We found an increased percentage of sperm containing DNA damage, as revealed by “comet tails” in *Aire^−/−^* samples compared to controls (**Fig. 7**; *Aire^+/+^*, 2.35 ± 0.92% vs. *Aire^−/−^*, 7.98 ± 3.69).

**Figure 7.**
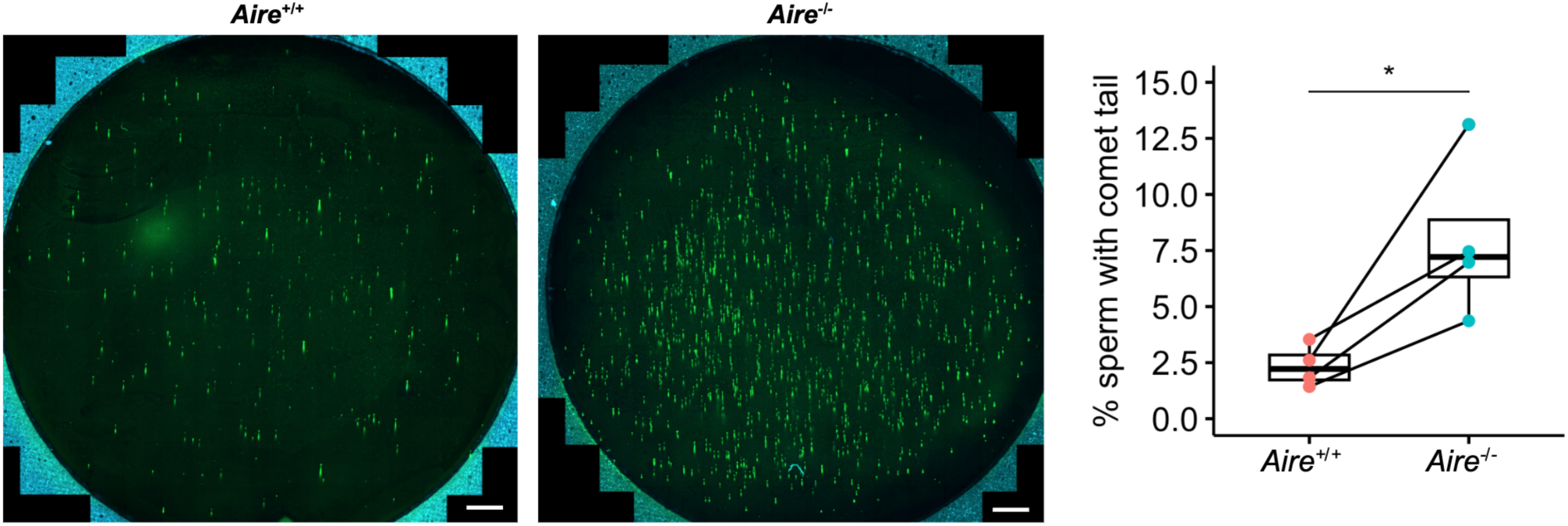
Comet assay reveal increased DNA fragmentation in caudal sperm from *Aire^−/−^* males. Epididymal tail sperm from *Aire^+/+^* or *Aire^+/-^* and *Aire^−/−^* males aged between 7 to 20 weeks were processed for comet analysis. Scale bar = 1000μm \**p* = 0.048, paired t-test.

### Decreased concentrations of Cu, Zn, and Se also correlate with increased sperm DNA fragmentation in *Aire^−/−^* animals

Oxidative stress due to increased reactive oxygen species (ROS) can result in DNA damage in cells.^41,42^ Because sperm are transcriptionally inert, they do not possess the ability for *de novo* synthesis of antioxidants and thus are especially prone to oxidative damage. To protect the sperm from such assault, the epididymal epithelial cells secrete antioxidant enzymes – glutathione peroxidases (Gpxs) and peroxiredoxins - into the luminal compartment^12,19,43^ for which essential elements such as Cu, Zn, and Se are of core structural component. To test this, we measured the concentration of metals from the caput, corpus, and cauda epididymis using ICP-QQQ-MS. Compared to *Aire^+/+^* and *Aire^−/−^;Rag1^−/−^*samples, concentrations of Cu, Zu, and Se were reduced, particularly in cauda epididymis of *Aire^−/−^* animals **(Supplemental Fig. 2, A-E).** Elemental analysis by LA-ICP-TOF-MS corroborated the findings for ^63^Cu and ^64^Zn, which were strikingly reduced in *Aire^−/−^* epididymis as compared to that of *Aire^+/+^*controls.

## Discussion

Male-factor infertility, which accounts for about half of these cases, can stem from several problems, including underlying autoimmune disease. In this instance, autoreactive lymphocytes may target the generative sources of male germ cells – the testis and epididymis – perturbing sperm maturation and negatively impacting fertility. We previously showed that male mice deficient in *Aire*, the transcription factor critical for establishing self-antigen specific tolerance to developing T cells in the thymus, suffered from subfertility with increased infiltration of CD3+ T cells into the testis and generation of anti-testis and anti-epididymal antibodies.^10^ Here, we report increased numbers of activated T cells and macrophages within the epididymis, which was associated with disruptions in inorganic physiology and sperm DNA damage.

We derived an *Aire*-deficient mouse model by targeting exon 2 of *Aire* using CRISPR/Cas9 technology. These mice recapitulated prior findings of *Aire* deficiency, in that both endocrine and non-endocrine organs showed evidence of autoimmunity as well as severe male subfertility. Importantly, the absence of adaptive immune cells, achieved by crossing Aire-deficient mice to *Rag1*-deficient mice, completely restored fertility. This observation conclusively demonstrates that infertility in *Aire^−/−^* males is driven by the presence of autoreactive lymphocytes rather than a germ cell or reproductive tract intrinsic role for Aire.

In many animals, we found high numbers of CD4+ and CD8+ T cells, as well as B cells, in the epididymis of *Aire^−/−^* males, while we did not observe testicular abnormalities. This result suggests that infertility may arise from impaired epididymal function. Using immunohistochemistry, we found that T cell abundance varied between animals but also showed as age- and region-specific differences in the epididymis. Additionally, flow cytometry revealed increased proportions of activated (CD69+) CD8+ T cells in all segments of the epididymis of *Aire^−/−^* animals. While CD44 expression was high on nearly all T cells regardless of genotype, the late activation marker, CD25, was upregulated in only CD4+ T cells from caudal *Aire^−/−^* epididymis, possibly indicating a memory phenotype in this region. Together, these data suggest that Aire deficiency promotes activation of CD8+ and CD4+ T cells, potentially due to chronic interaction between the T cell receptor and self-antigens presented in the context of MHC class I or class II in the Aire^−/−^ epididymal environment. Future experiments will include additional markers that will reveal memory and exhaustion phenotypes of these cells.

In addition to characterizing the immune cell infiltrates, we set out to understand the potential mechanism of fibrosis in autoimmune-mediated epididymitis by combining immunohistochemical staining and mass spectrometry mapping of the metal content using LA-ICP-TOF-MS and ICP-QQQ-MS. While epididymal fibrosis is a hallmark feature of inflamed epididymis due to infection or experimentally induced autoimmune epididymitis,^44,45^ our findings of interstitial iron overload as well as ferritin-positive interstitial macrophages are reminiscent of hemochromatosis and bleomycin-induced idiopathic pulmonary fibrosis where the associations between iron, macrophages, and fibrosis are well-established.^31,33,46–48^ It should be recognized that, although macrophages likely account for most F4/80+ cells, other cells express this marker (eg, a subset of dendritic cells^49^, and eosinophils^36^) and thus may also be responsible for carrying iron. Conversely, ferritin+/F480-negative cells may also account for excess iron in the Aire^−/−^ epididymides. Dysregulation of iron chemistry can promote oxidative damage through Fenton reaction and increase hydroxyl radicals that readily cause cellular DNA damage, which may instigate fragmentation of sperm DNA and fibrosis of the interstitium through the wound healing process. While the sequence of events – whether accumulation of iron promotes fibrosis such as that observed in the lung^48^ or vice versa^33^ – is unknown, we surmise that the autoimmune injury to the epididymis activates F4/80+ interstitial macrophages evidenced by their morphological changes, which by working together with myofibroblasts, promotes fibrosis of the epididymal interstitium.^50^

We show that autoimmune-mediated epididymitis decreases the concentrations of essential metals in the epididymis, including Cu, Zn, and Se. These findings were surprising because imbalance of these elements in seminal fluid is associated with sperm abnormality and infertility in humans,^51,52^ To protect the sperm from oxidative damage, the epididymal epithelial cells express a wide variety of antioxidant enzymes including Cu-Zn superoxide dismutases (including SOD1 and SOD3^53^), glutathione peroxidases (Gpxs^11,54–56)^, and thioredoxin-glutathione reductase (i.e. TXNRD3^57^). These enzymes have been shown to be released into the lumen in the form of epididymosomes by the epithelial cells.^58^ Deficiency of these essential cofactors may confer deleterious effect on the activity of key antioxidant enzymes including SODs and selenoproteins TXNRD3, Gpx1, 3, and 4, exposing sperm to reactive oxygen species-mediated oxidative stress and DNA damage.^59,60^ We speculate that decreases in Cu, Zn, and Se lead to defects in antioxidant enzyme production or function, resulting in sperm DNA damage. While further studies are required to confirm levels of antioxidant activity and production of reactive oxygen species, this model provides a plausible explanation for why sperm from *Aire^−/−^* males fail to allow embryo progression to blastocyst stage upon *in vitro* fertilization.^60,61^

In summary, we demonstrate that Aire deficiency and resultant autoimmune attack to the epididymis disrupts inorganic physiology of the epididymis, in particular accumulation of Fe in the interstitial space, and decreases in Zn, Se, Cu, which may in turn promote increased ROS production and damage to sperm DNA. Interstitial macrophages as well as myofibroblasts likely play a significant role in the fibrogenesis using similar mechanisms found in the liver and the lung.^62^ Based on the results from *Aire^−/−^;Rag1^−/−^* animals, we demonstrate that fibrogenesis, Fe accumulation, and Zn, Cu and Se depletion in the interstitium require the presence of adaptive immune cells. Future studies will focus on further characterizing these immune population in the *Aire^−/−^*epididymis to delineate how they participate in the fibrogenic transformation of the epididymis and promote imbalance of essential metals that are critical to maintain male fertility. Further studies are necessary to understand the mechanisms that contribute to decreases in essential metals such as Se, Cu, and Zn in autoimmune-mediated subfertility.

## Acknowledgements

We thank Dr. Lindsay Moritz and Dr. Sue Hammoud at University of Michigan for the provision of the comet assay protocol. Metal analysis using ICP-QQQ-MS and LA-ICP-TOF-MS was performed at the Quantitative Bio Element Analysis and Mapping Center (QBEAM) located at Michigan State University, East Lansing, MI, USA. Figures were created using BioRender.

---

**Supplementary Figure 1.**
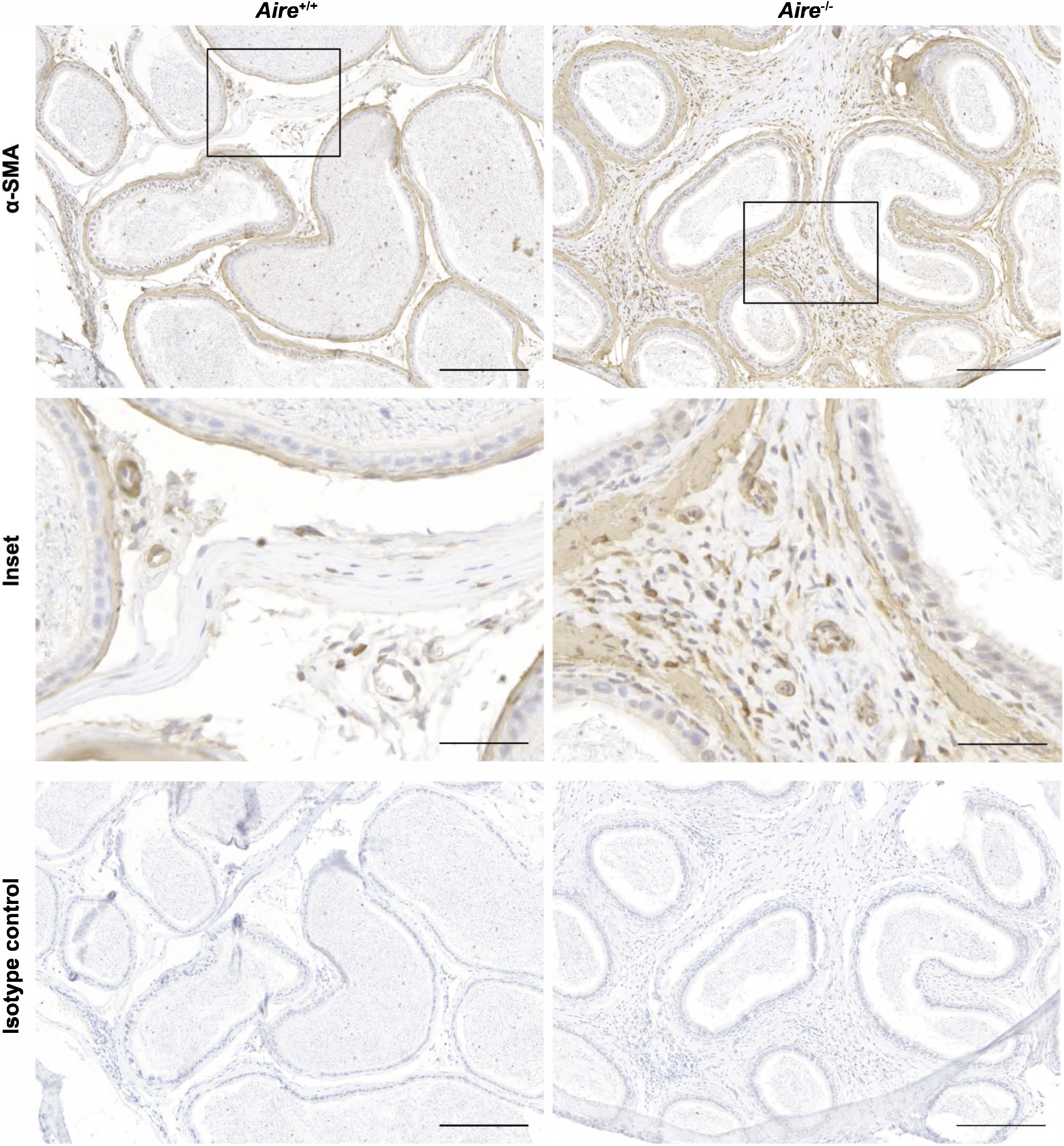
αSMA+ myofibroblasts abound in the interstitium of *Aire^−/−^*cauda epididymis in comparison to *Aire^+/+^* epididymis of similar age. Scale bar = 200 μm for top and bottom row; 50 μm for middle row

**Supplementary Figure 2.**
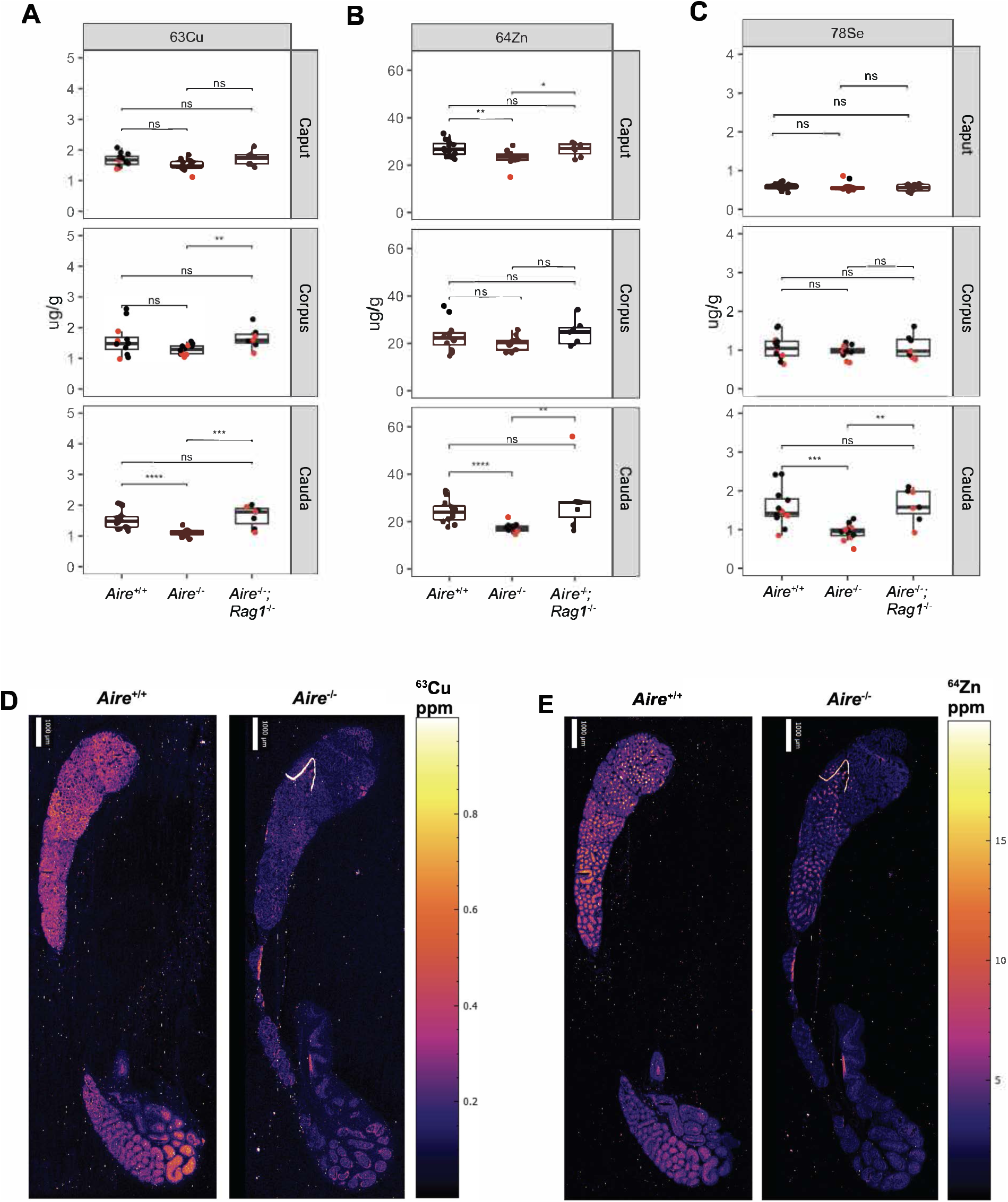
Concentrations of ^63^Cu and ^64^Zn are decreased in *Aire^−/−^* epididymis. **(A-C)** ICP-QQQ-MS analysis of individual segments of the epididymis show regional decreases in ^63^Cu, ^64^Zn, and ^78^Se in the *Aire^−/−^* epididymis compared to *Aire^+/+^* or *Aire^−/−^;Rag1^−/−^*. Red = 7-9 weeks old, Black = 19-20 weeks old. Elemental distribution of ^63^Cu Statistical analysis: Kruskal-Wallis One-Way ANOVA followed by Dunn-Bonferroni post-hoc was conducted to compare the means of each group. *p**(D)** and ^64^Zn **(E)** in 20-week-old *Aire^+/+^* and *Aire^−/^*^−^ epididymis using LA-ICP-TOF-MS; Se abundance was below the detection limit of this technology. Scale bar = 1000 μm.

**Supplementary Table 1.** Elemental concentration in the gelatin standards.

| **Sample Name** | **<sup>24</sup>Mg** | **<sup>44</sup>Ca** | **<sup>55</sup>Mn** | **<sup>57</sup>Fe** | **<sup>58</sup>Ni** | **<sup>63</sup>Cu** | **<sup>64</sup>Zn** |
| --- | --- | --- | --- | --- | --- | --- | --- |
| **Gelatin Blank** | 2.106 | 0.916 | 0 | 0.679 | 0.013 | 0.030 | 0.096 |
| **Gelatin 50 ppm** | 53.32 | 44.85 | 55.40 | 56.99 | 56.55 | 56.93 | 55.02 |
| **Gelatin 20 ppm** | 26.62 | 21.17 | 26.39 | 26.97 | 27.03 | 27.64 | 26.34 |
| **Gelatin 10 ppm** | 14.38 | 11.01 | 13.66 | 14.14 | 14.01 | 14.13 | 13.08 |
| **Gelatin 5 ppm** | 8.512 | 6.498 | 7.535 | 8.074 | 7.409 | 7.821 | 7.243 |
| **Gelatin 1 ppm** | 3.460 | 2.113 | 1.681 | 2.564 | 1.768 | 1.832 | 1.776 |

**Supplementary Table 2.** LA-ICP-TOF-MS parameters.

| ***ICP-MS Paramters (Tofwerk S2)*** |  |  |
| --- | --- | --- |
| **Parameter** | **Unit** | **Value** |
| RF Power | W | 1550 |
| Sampling Depth | mm | 4.9 |
| Cone Material | Nickel | - |
| Cone Insert (STD) | mm | 3.5 |
| Plasma Gas Flow | L/min | 14.0 |
| Auxillary Gas Flow | L/min | 8.0 |
| Nebulizer Gas Flow | L/min | 0.95-1.01 |
| Measurement Mode | CCT Mode | - |
| CCT Gas Flow (100% He) | mL/min | 5 |
| CCT Focus lens | V | -19.5 |
| CCT Entry Lens | V | -180 |
| CCT Mass | V | 250 |
| CCT Bias | V | 10 |
| CCT Exit Lens | V | -200 |
| ***Time-of-Flight Parameters (Tofwerk S2)*** |  |  |
| **Parameter** | **Unit** | **Value** |
| m/z range | amu | 14 - 256 |
| Resolution | m/Δm | 1000 |
| ODG Settings | ms | 24 |
| Notch Bias | V | -80 |
| Notch (40 amu) | V | 1.6 |
| Notch (36 amu) | V | 1 |
| Notch (28 amu) | V | 2 |
| Notch (15.8 amu) | V | 2 |
| ***Laser Ablation Parameters (ESL Bioimage 266 nm)*** |  |  |
| **Parameter** | **Unit** | **Value** |
| Spot Size | μm | 10 |
| Interline Distance (y) | μm | 10 |
| Overlap (x) | μm | 0 |
| Repetition Rate | Hz | 100 |
| Laser Power | % | 60 |
| Laser Fluence | J/cm<sup>2</sup> | 15 |
| Sample Energy | mJ | 0.010-0.015 |
| Imaging Cup Flow Rate (He) | mL/min | 200 |
| Imaging Chamber Flow Rate (He) | mL/min | 250 |
| PEEK Tubing I.D. | mm | 0.75 |

## Notes

**Funding:** This research is supported by NIH grants R01 HD100832, R01 GM11484 and the National Research Resource for Quantitative Mapping in the life Sciences, supported by the Office of the Director, National Institutes of Health, and the National Institute for General Medical Sciences under grant number P41 GM135018.

### Competing Interest Statement

The authors have declared no competing interest.

### Summary of Updates

The previous manuscript version included formatting error that we wanted to address.

